# Hydrothermal Vent Species Assemblage Networks Identify Regional Connectivity Patterns in the Northwest Pacific

**DOI:** 10.1101/2022.07.20.500901

**Authors:** Otis Brunner, Chong Chen, Thomas Giguere, Shinsuke Kawagucci, Verena Tunnicliffe, Hiromi Watanabe, Satoshi Mitarai

## Abstract

The distribution of species among spatially isolated habitat patches supports regional biodiversity and stability, so understanding the underlying processes and structure is a key target of conservation. Although multivariate statistics can infer the connectivity processes driving species distribution, such as dispersal and habitat suitability, they rarely explore structure. Methods from graph theory, applied to distribution data, give insights into both connectivity pathways and processes by intuitively formatting the data as a network of habitat patches. We apply these methods to empirical data from the hydrothermal vent habitats of the Northwest Pacific. Hydrothermal vents are ‘oases’ of biological productivity and endemicity on the seafloor that are imminently threatened by anthropogenic disturbances with unknown consequences to biodiversity. Here, we describe the structure of hydrothermal vent species assemblage networks, how local and regional parameters affect their structure, and the implications this has for conservation. Two complementary networks were formed from an extensive species assemblage dataset: a bipartite network of species nodes linked to vent site nodes at which they are present, and a similarity network of vent site nodes linked by weighted edges based on their pairwise assemblage similarity. Using these networks, we assessed the role of individual vent sites in linking their network and identified biogeographic sub-regions. The three sub-regions and two outlying sites are separated by their spatial arrangement and local environmental filters. Both networks detected vent sites that play a disproportionately important role in regional pathways, while the bipartite network also identified key vent sites maintaining the distinct species assemblages of their sub-regions. These regional connectivity pathways provide insights into historical colonisation routes, while sub-regional connectivity pathways are of value when selecting sites for conservation and/or estimating the multi-vent impacts from proposed deep-sea mining.

## Introduction

Conservation efforts aim to slow the global degradation of biodiversity (Meine et al. 2006) as well as ecosystem functions and services (Nicholson et al. 2009). These features of any ecosystem are supported by ecological connectivity, the flow of organisms, energy, and materials across suitable habitat patches (Crooks and Sanjayan 2006; Correa Ayram et al. 2016). Structural or functional isolation of habitat patches by natural or anthropogenic disturbances may limit the capacity of an ecosystem to maintain processes that are valued within conservation objectives (Rudnick et al. 2012) by disrupting landscape connectivity. If the dispersal of individuals is impeded, the entire species may become more vulnerable to global extinction by reducing their ability to shift their range in response to climate change or support local recovery following disturbance events (reviewed by Jones *et al*., 2007). For these reasons, maintaining the structure of landscape connectivity is a global priority (IUCN 2017). Island or ‘island-like’ systems (*sensu* (Dawson and Santos 2016)) are particularly vulnerable to disturbances because of their characteristically isolated nature (Wilson and MacArthur 1967; Losos and Ricklefs 2009). Hydrothermal vents are island-like ecosystems (Dawson and Santos 2016; Mullineaux et al. 2018) that exhibit particularly high levels of biomass and endemicity at the seafloor (Corliss et al. 1979; Van Dover 2014).

Hydrothermal vents are often described as ‘oases’ of high biomass in the deep (Laubier 1993), as local chemoautotrophy not only supports higher densities of benthic species within the vent ecosystem, but also contributes to the surrounding non-vent ecosystems (reviewed in (Levin et al. 2016). Vents are also home to seafloor massive sulfide deposits, a primary target for deep-sea mining in recent years (Van Dover et al. 2018). Where these ecosystems coincide with mining interests, regional diversity may become vulnerable (C. L. Van Dover et al. 2018) and a global priority for protection (Thomas, Böhm, et al. 2021). In the Northwest Pacific, where the first test mining of hydrothermal vents occurred in the Okinawa Trough (Okamoto et al. 2019), studies have assessed the impact on the local vent community and surrounding habitats at the site of proposed mining (Nakajima et al. 2015). Although over three-fourths of vent endemic species in the Northwest Pacific are considered threatened by deep-sea mining (Thomas et al. 2021a), very few studies have attempted to assess the effect this proposed mining will have on the regional vent communities (Suzuki et al. 2018). Here we use empirical observations to describe the structure of connectivity among spatially isolated vent communities to investigate the regional impacts of deep-sea mining.

Connectivity among vent communities is facilitated by the dispersal of planktonic larvae (Adams et al. 2012). Although it is possible to quantify dispersal probabilities using oceanographic simulations (Mitarai et al. 2016), dispersal is just one of several processes required for demographic connectivity among discrete communities. Reproduction, larval dispersal, settlement, and maturation are the sequential steps necessary to maintain demographic connectivity among sites (Kritzer and Sale 2004). Therefore, connectivity is controlled by a combination of local and regional processes such as dispersal probability, habitat suitability and biological interactions (reviewed in (Pineda et al. 2007)). Here we investigate the drivers of diversity by formatting species’ distributions as a network and then test how local and regional drivers of diversity can explain the structure of this network. Such ‘similarity networks’ have seldom been applied to empirical observations of linked but spatially distinct communities (metacommunities) is limited (Borthagaray et al. 2015), despite a growing body of literature has theoretically demonstrating the value of metacommunity networks to the study of biodiversity in general (Keitt et al. 1997; Economo and Keitt 2010; Suzuki and Economo 2021).

Previous studies have also presented hydrothermal vents as networks and applied methods from graph theory to detect biogeographic regions and infer historical connectivity pathways at the global scale (Moalic et al. 2012; Kiel 2016). Here, we focus on similar pathways among hydrothermal vent sites of the Northwest Pacific at an intra-regional scale more relevant to contemporary conservation. The Northwest Pacific is a distinct biogeographic region in terms of vent-endemic fauna (Bachraty et al. 2009) and in terms of dispersal through oceanographic simulations (Mitarai et al. 2016). Within this region, the interactions between the Pacific Plate, the Philippine Plate and the Eurasian Plate create active tectonic margins containing trenches, volcanic arcs and back-arc spreading centres. Hydrothermal vents are present in the volcanic arcs and back-arcs of the Izu-Bonin, Mariana and Okinawa basins (Figures 1). In these arc-back-arc basins, 78 known hydrothermal vent sites are recognized by the InterRidge database (Beaulieu and Szafranski, 2020). Although the Northwest Pacific is one of the better surveyed regions in terms of hydrothermal vent biodiversity, few regions are considered to have been surveyed comprehensively (Thaler and Amon 2019). By considering the vent sites of the Northwest Pacific as an interconnected network, we can apply structural analyses from graph theory to assess the roles individual vent sites play in sharing species across the region. Maintaining connectivity through shared species within a hydrothermal vent biogeographic region is a conservation priority due to its importance to biodiversity and its vulnerability to anthropogenic disturbances (Turner *et al*., 2019). We use empirical observations to investigate the processes that maintain connectivity - such empirical-based observations have been identified as an important knowledge gap (Van Dover 2014, Amon *et al*., 2022)

**Figure 1:**
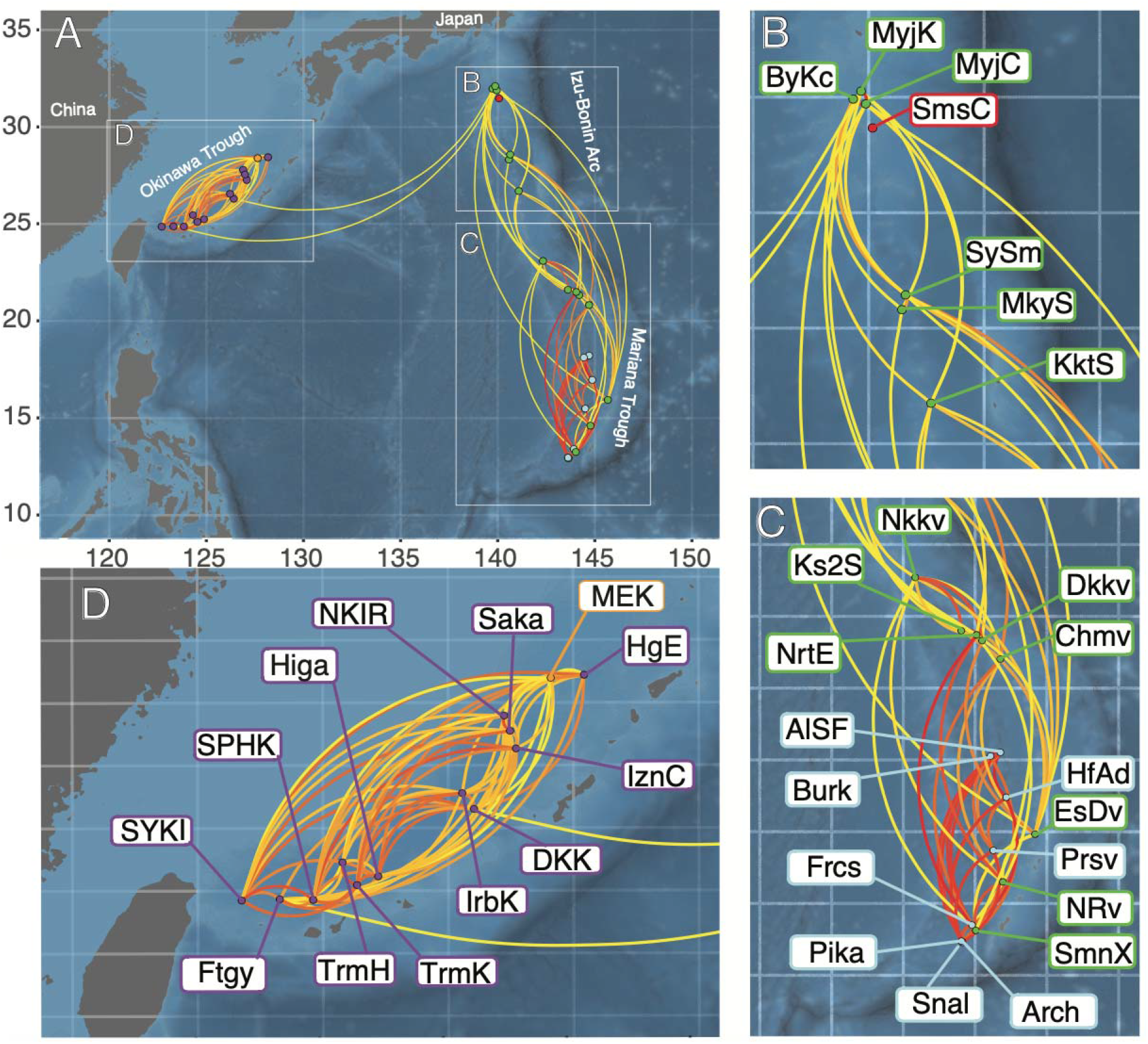
The vent sites of: (a) the Northwest Pacific, (b) Izu-Bonin, (c) Mariana and (d) Okinawa that were used in this study. The colour of the vent sites represent the distinct sub-regions as detected from the Modularity analysis (purple – Okinawa Trough (OT), orange – Minami-Ensei Knoll, red – Sumisu, green – Izu-Bonin-Mariana Arc, blue – Mariana Trough (MT)). Lines connecting vent sites represent the pairwise similarity (Sorensen’s Coefficient) between species assemblages at vent sites. Red lines represent higher similarity while yellow lines represent lower similarity.

Hydrothermal vent species occurrence data have previously been used to detect biogeographic barriers within the Northwest Pacific (Nakajima et al. 2014; Kojima and Watanabe 2015; Watanabe and Kojima 2015; Watanabe et al. 2019; Giguère and Tunnicliffe 2021). These studies have inferred connections using the number of shared species or β-diversity between sites and/or SIMPROF analysis (Clarke et al. 2008) to detect significantly similar species assemblages among vent sites, and infer the biogeographic barriers that separate others. In this study, we combined and curated the occurrence data from these previous studies along with new occurrence data to create a comprehensive view of the regional species assemblages, with representation from the three major arc-back-arc systems of the region. For comparability, we first replicated the same clustering methods used in the aforementioned studies. We then expanded and improved upon the previous studies by applying methods from graph theory, which offers distinct advantage over more classical pairwise analyses of connectivity (Proulx, Promislow, and Phillips 2005) and the detection of biogeographic barriers (Bloomfield, Knerr, and Encinas-Viso 2018). As the connectivity patterns of the regional networks are inferred from shared species, they provide insights into the geological and biogeographic history of the region and the ability to identify those vent sites central to regional species distribution patterns.

## Methods

### Species Occurrence Data

We assembled occurrence records of vent-associated benthic megafauna from a variety of published and new data. The occurrence records were identified to the lowest taxonomic level with some published records being updated based on a review of recent taxonomic literature. Only species-level records were used in this study to ensure compatibility between the different data sources and because this is the required taxonomic resolution when studying processes at the metacommunity and sub-regional scale (Webb et al. 2003). Species names were checked against the World Register of Marine Species (WoRMS, http://www.marinespecies.org/) to ensure up-to-date nomenclature. All occurrences were associated with a named vent site in the InterRidge database (Beaulieu and Szafranski, 2020) based on its geographic location or associated metadata to create a site-by-species matrix of 36 vent fields and 117 species (Supplemental Table 1). Due to the remote nature of vent ecosystems and the difficulty in carrying out comprehensive surveys, this matrix is a ‘presence-only’ dataset, as species absence from vent sites cannot be confirmed.

### Similarity Network

We calculated the Sørensen’s coefficient (Sorensen 1948) from the site-by-species matrix to give a pairwise dissimilarity between all vent sites based on their species assemblages. Sørensen’s coefficient was used as it is applicable to presence-absence data sets, but gives extra weight to shared presence (Legendre and Legendre 2012). Furthermore, Sørensen’s coefficient was used to compare results with those of previous studies in this region (Nakajima et al. 2014; Kojima and Watanabe 2015; Watanabe and Kojima 2015; Watanabe et al. 2019) which also used this coefficient. A subsequent SIMPROF analysis (Clarke, Somerfield, and Gorley 2008) used 1000 permutations at 5% significance to hierarchically group vent sites into clusters (hereafter referred to as SIMPROF clusters to avoid confusion with the defined term “network cluster” used in graph theory) that had similar species assemblages.

Using the pairwise similarity (1 – Sørensen’s coefficient) between vent sites, we created a network of vent sites in the Northwest Pacific. The similarity value was used as the weight of the edges that link the vent site nodes in this network, hereafter referred to as a ‘similarity network’. The ‘percolation threshold’ of the similarity network was calculated following the methods of Rozenfeld *et al*., (2008) using the ‘sidier’ package in R (Muñoz-Pajares, 2013; R Core Team, 2021) and used to remove ‘weak’ links. The relative importance of each vent in maintaining a connected network was then evaluated based on their ‘betweenness centrality’ (Freeman 1977), the frequency they occur in the geodesic path between each pair of vents in the network. Betweenness centrality was calculated for every node in the similarity network after thresholding using the ‘igraph’ package in R (Csardi and Nepusz 2006).

We used variance partitioning to determine the contribution of environmental and spatial parameters to explain the β-diversity (Legendre and Legendre 2012) represented by the edge weight between nodes in the species assemblage network. The spatial parameter in question was calculated using ‘distance-based Moran’s Eigenvector Maps’ (dbMEM) following the methods detailed in Legendre and Legendre (2012). These dbMEMs summarise the relative position of each site based on their geodesic distance from their neighbouring sites. Two sets of dbMEM were created; the ‘fine-scale dbMEM’ uses a threshold of geodesic distance lower than that required to connect sites within the Okinawa Trough to other sites in the Northwest Pacific, while the ‘broad-scale dbMEM’ does consider this connectivity when calculating relative position. The environmental parameters tested were the depth and tectonic setting as recorded in the InterRidge database (Beaulieu and Szafranski, 2020) with some additional corrections (TABLE 1). These local environmental variables were selected because they have been recorded for every vent site in the dataset and are indicative of the many processes that directly affect local habitat suitability (Tunnicliffe, McArthur, and McHugh 1998; Mullineaux et al. 2018; Giguère and Tunnicliffe 2021). The variance partitioning analyses as well as the formation of the dbMEMs were carried out using the ‘vegan’ package in R (Oksanen et al. 2019)

**Table 1:**
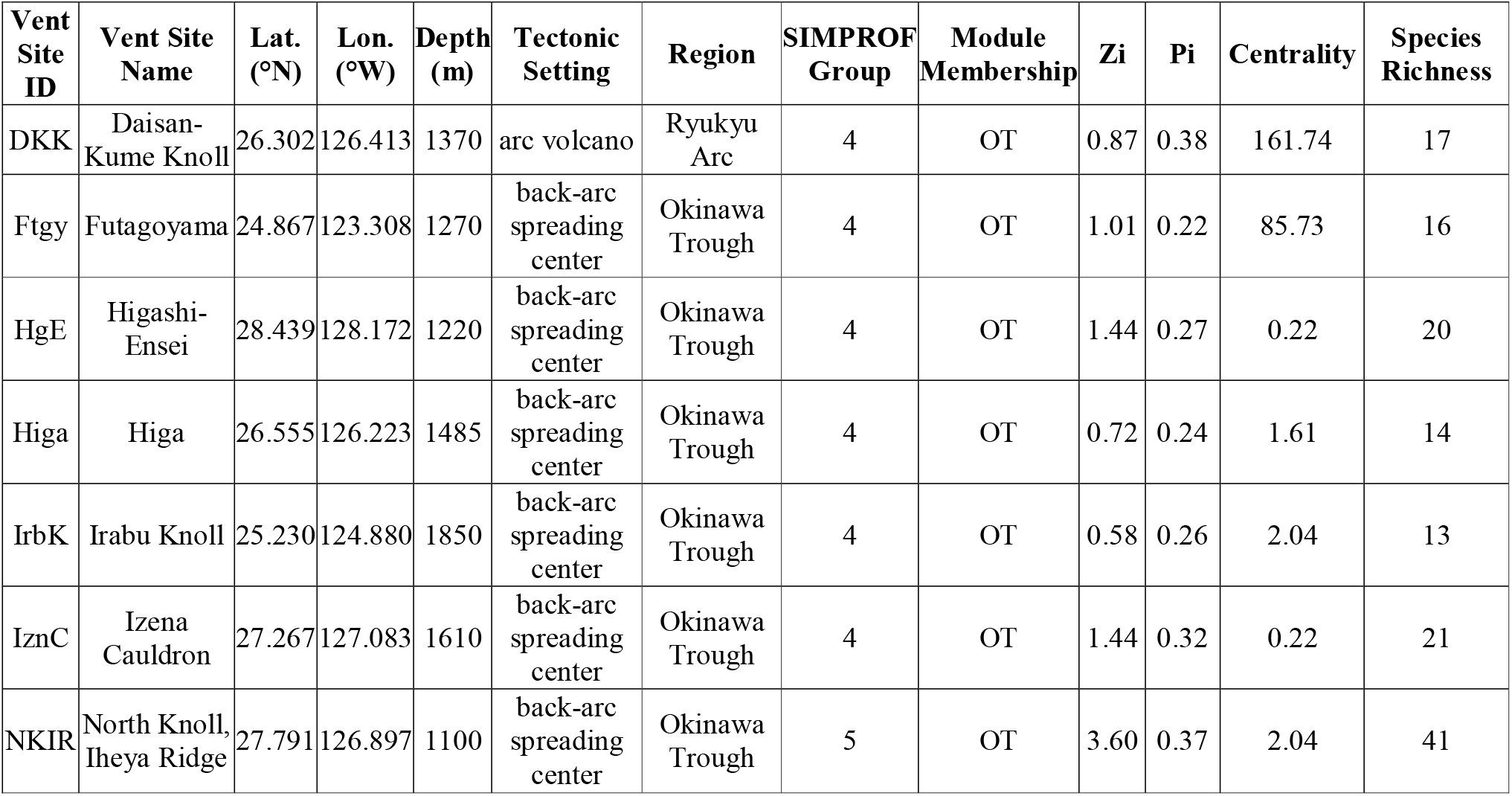
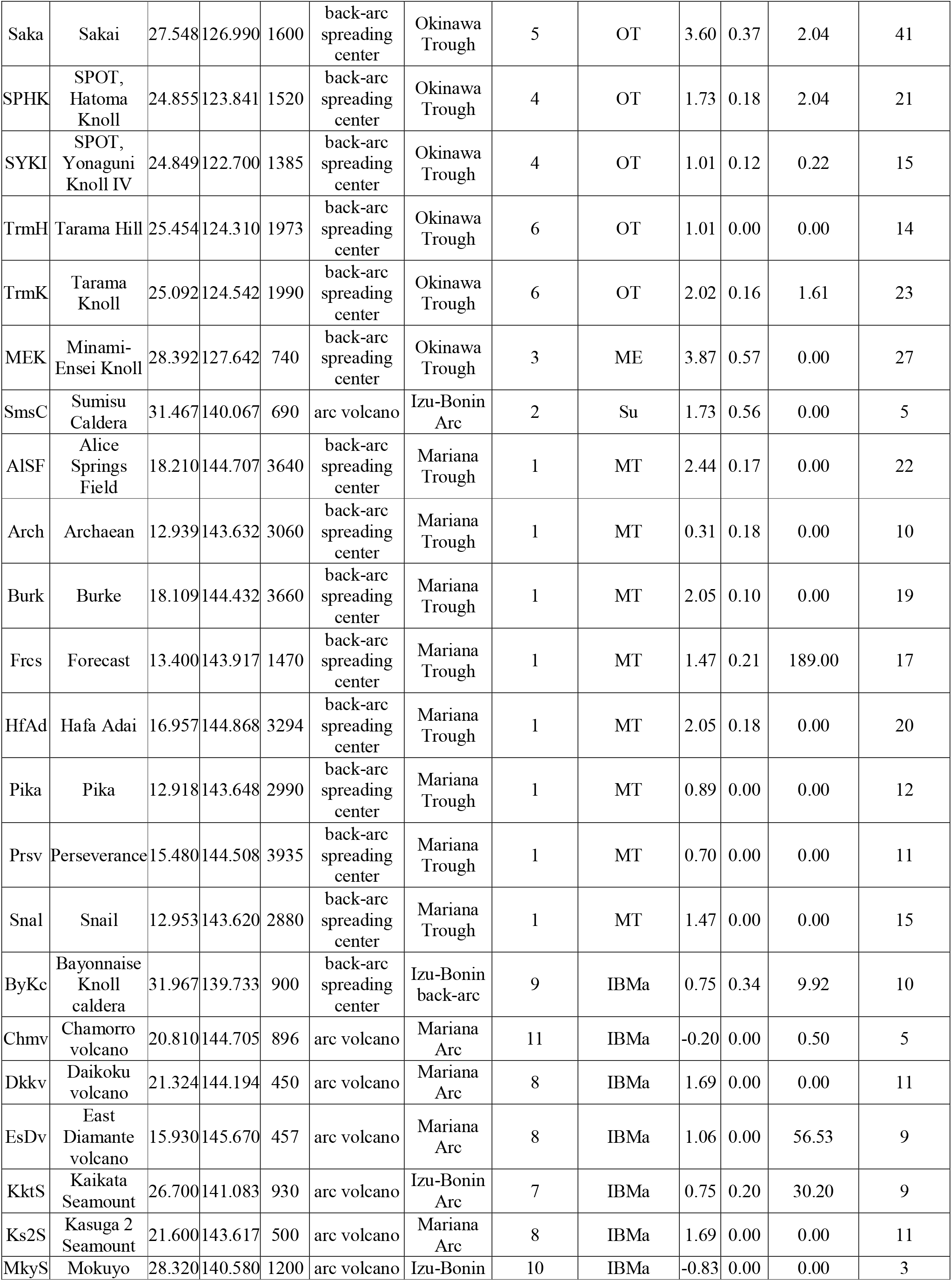
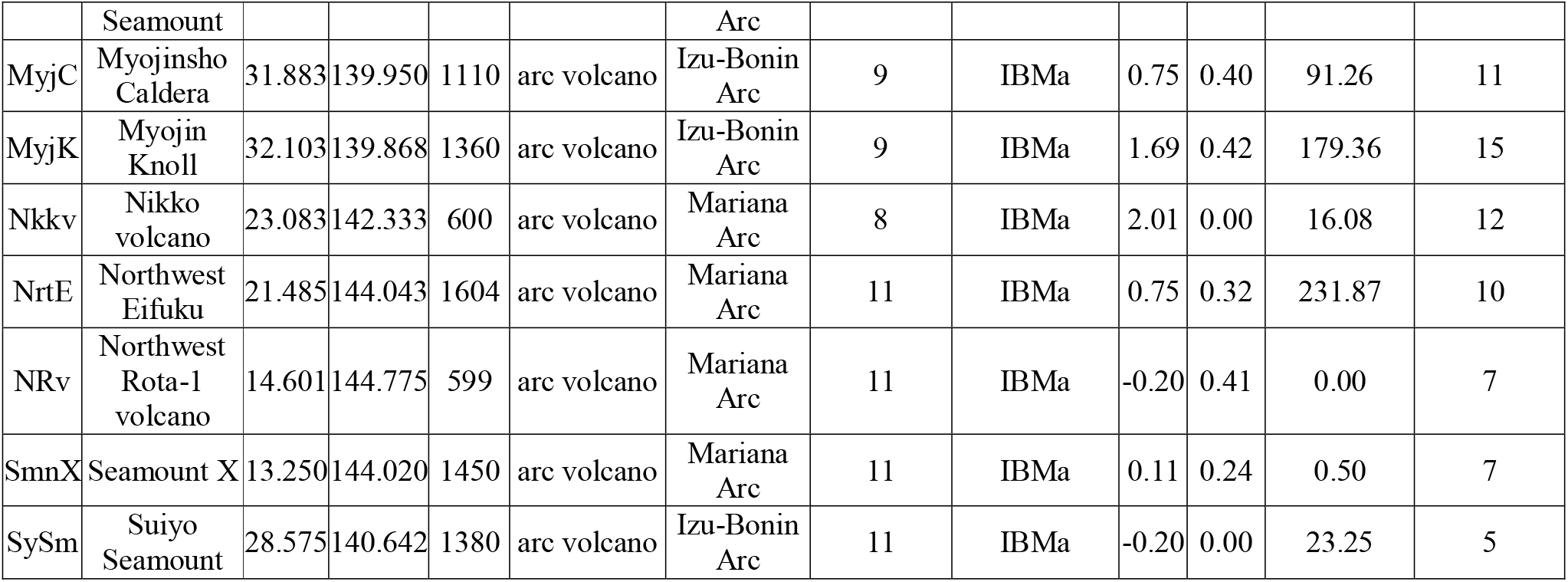
Description of vent sites used in analyses

### Bipartite Network

The second network was formed directly from the site-by-species matrix and is referred to as a bipartite network, after the two types of nodes it contains. The first node type, a species node, is linked to the second type, a vent site node, if the species was present at said vent site. The nodes in this network are linked by unweighted edges which only occur between nodes of a different type (i.e. species – vent site). This network approach has been implemented in various biogeographic studies (Carstensen et al. 2012; Dalsgaard et al. 2014; Kougioumoutzis et al. 2014; Kougioumoutzis et al. 2017) to detect barriers to ‘biogeographical connectivity’. Following the methods of Cartensen *et al*. (2012), a simulated annealing approach was used to subdivide the regional bipartite network iteratively into groups until the grouping that maximises the ‘Modularity’ value of the network is identified. The ‘Modularity’ is a measure of the extent to which nodes have more links within their group than expected if the links are random (Guimerà and Nunes Amaral 2005; Guimerà and Amaral 2005). For this analysis we used the ‘rnetcarto’ package in R (Doulcier and Stouffer 2015; R Core Team, 2021). The resultant groups of highly linked nodes in the bipartite network are hereafter referred to as ‘modules’. A redundancy analysis was used to detect the possible roles of known biogeographic barriers – depth, tectonic setting, and distance (broad-scale dbMEM) – on module membership, following the recommendations of Legendre and Legendre (2012). Additionally, a MANOVA analysis was carried out with the same formula to detect any significant variation of each explanatory variable and their interacting terms between the module groups.

Each node’s role in connecting the bipartite network was assessed based on its within-module degree (z_i_) and participation coefficient (P_i_) (Guimerà and Nunes Amaral 2005; Guimerà and Amaral 2005). As the direct links to a vent site node come from the species it contains, the position of a vent site node in z_i_ - P_i_ space is indicative of its species richness, the regional distribution of those species and the role the site itself plays in connecting spatially isolated species assemblages (Carstensen et al. 2012). Each vent node was assigned one of the universal cartographic roles defined by Guimera and Amaral’s (2005b). These roles are as follows: ‘Peripheral nodes’ (z_i_ < 2.5 and P_i_ < 0.62), ‘Module hubs’ (z_i_ > 2.5 and P_i_ < 0.62), ‘Connector nodes’ (z_i_ < 2.5 and P_i_ > 0.62), and ‘Network Hubs’ (z_i_ > 2.5 and P_i_ > 0.62). These metrics of module connectivity were calculated using the ‘rnetcarto’ package in R (Doulcier and Stouffer 2015).

## Results

### Similarity Network

The similarity network shows three distinct groups in the form of qualitative network clusters (Newman 2010) once the percolation threshold of 0.7 was applied. The edges that remain after this threshold and how they connect vent sites can be seen in a geographic (Figure 1) or simplified layout (Figure 2). The simplified layout positions vent sites relative to others with which they are directly linked, revealing three qualitative network clusters. These three network clusters are the vent sites of the Mariana Back-arc, the Okinawa Trough, and the Izu-Bonin-Mariana Arc. The betweenness centrality of nodes (Table 1) was high for those vent sites that linked the three network clusters: Northwest Eifuku, Forecast and Myojin Knoll. However, the SIMPROF analysis returned eleven groups (SIMPROF clusters) of vent sites each of which has no significant structural differences among their species assemblages (Figure 3).

**Figure 2:**
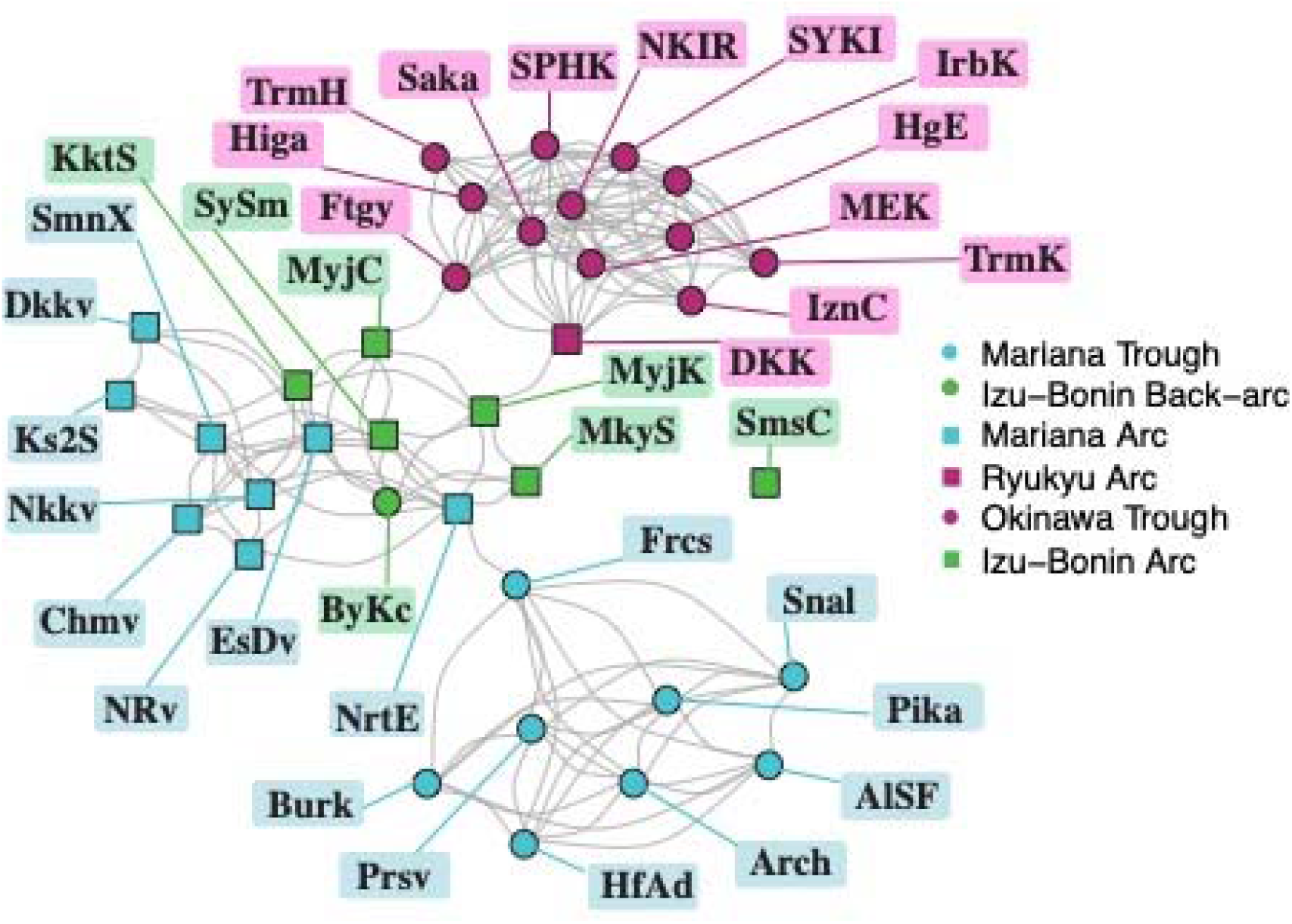
Similarity network of the Northwest Pacific vents showing qualitative clustering based on tectonic basin. The shape represents the tectonic setting of each vent site and colour represents their basin. Edges represent pairwise similarity values between vent nodes above the percolation threshold. The relative position of the vent nodes is dictated by the shared edges.

**Figure 3:**
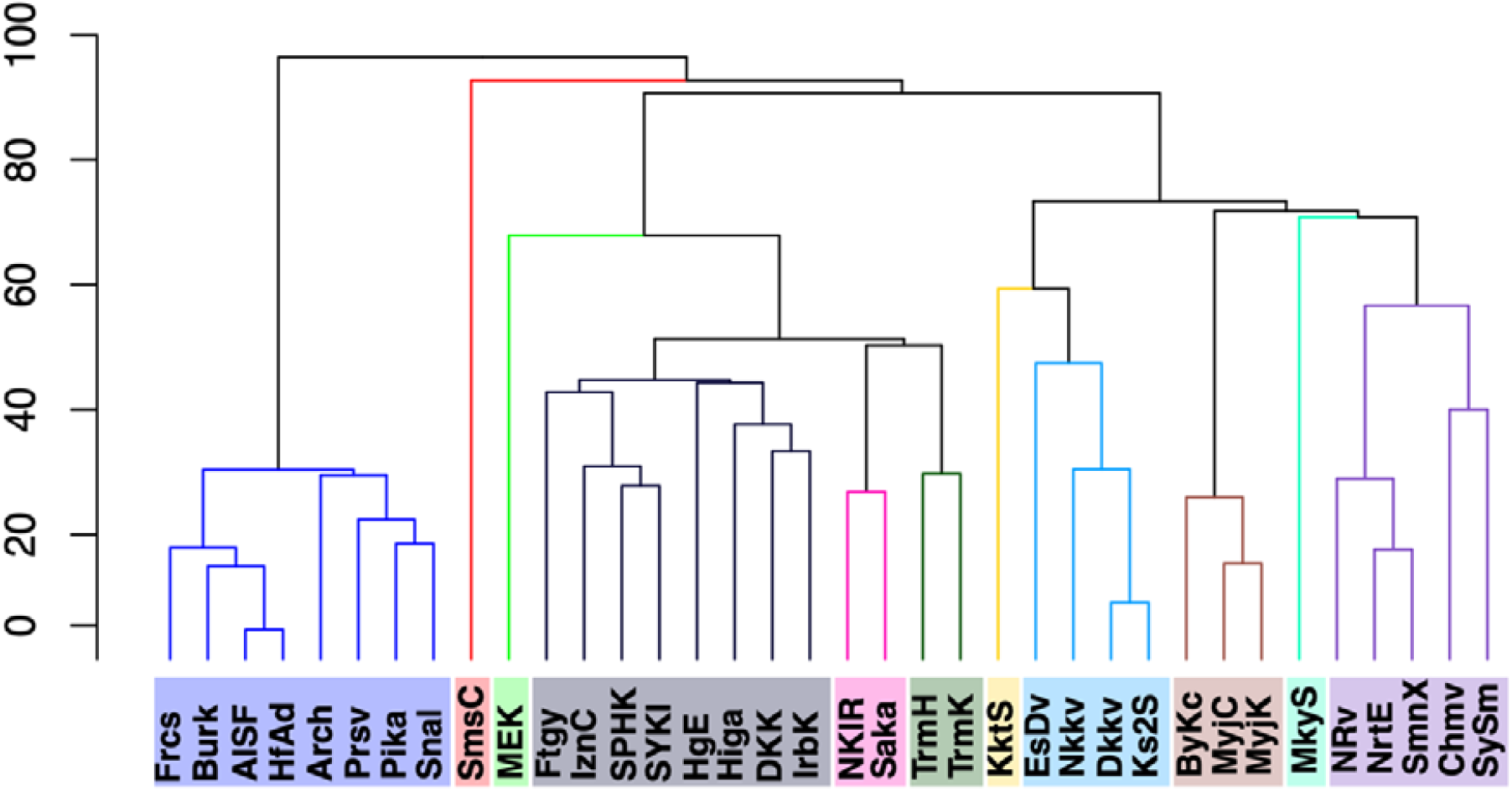
SIMPROF clustering of Northwest Pacific vent sites based on Sorensen’s coefficient

The explanatory variables used in the variance partitioning (Figure 4) explain 80% of the variation in the Sørensen’s Coefficient between vent sites. Much of this variation is explained by the broad-scale dbMEM; alone, it explains 22% while its interaction with the environmental parameters of depth and tectonic setting explains an additional 25%. Alone the environmental variation between vent sites explains 10% and a further 4% when combined with the fine-scale dbMEM spatial parameter. The fine-scale dbMEM only explains 7% of variation on its own and a further 15% as an interaction with the broad-scale dbMEM. Depth was checked for spatial autocorrelation following the methods outlined in Legendre and Legendre (2012) and found to be non-significant.

**Figure 4:**
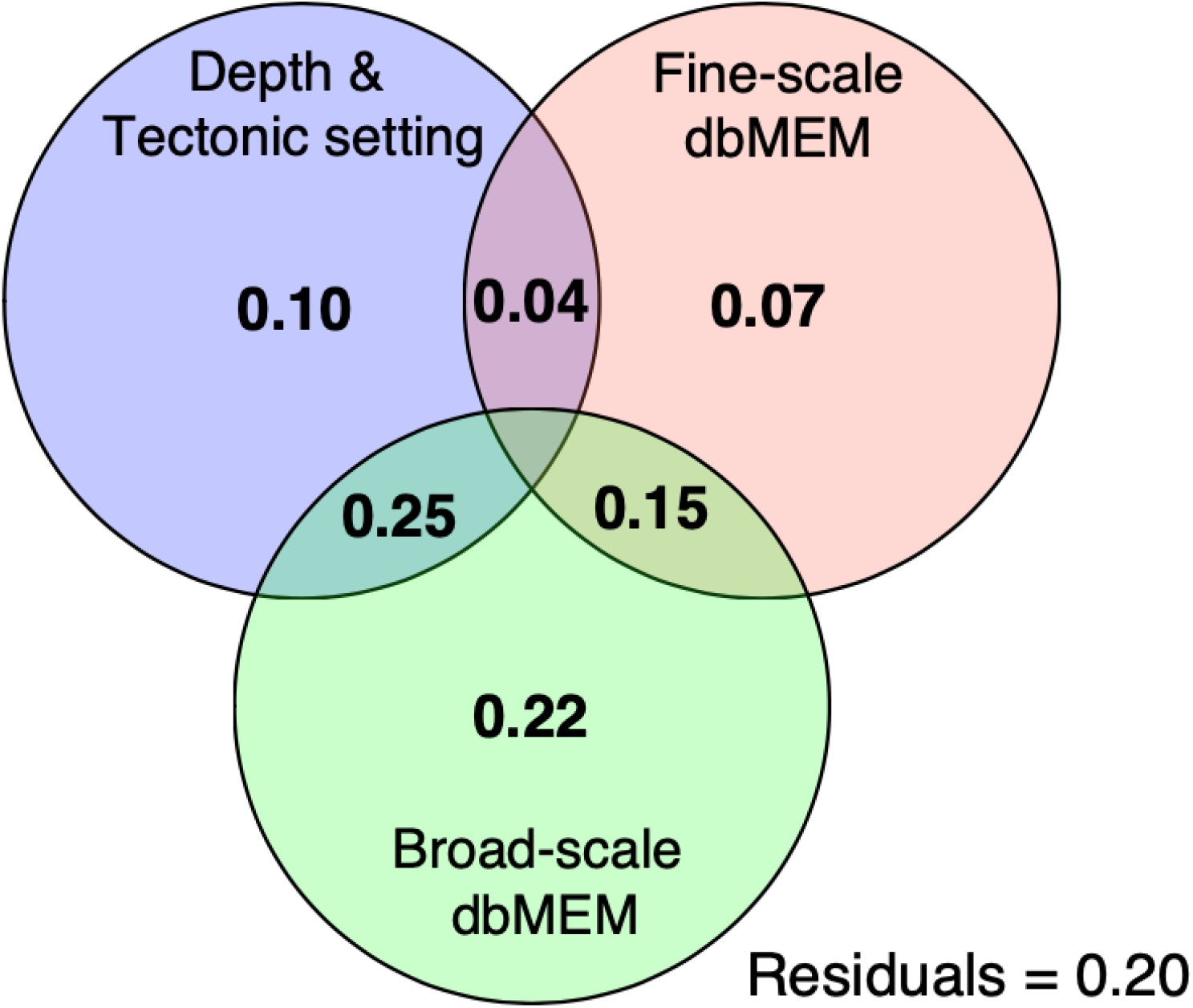
Variance Partitioning of vent site dissimilarity (Sorensen’s coefficient) against local environmental variables (depth and tectonic setting), fine-scale dbMEM and broad-scale dbMEM. The low residuals show how well these variables predict dissimilarity, particularly the Broad-scale dbMEM.

### Bipartite Network

The simulated annealing method (Guimerà and Nunes Amaral 2005; Guimerà and Amaral 2005) detected five distinct modules (Figure 5). Hereafter, we call these modules: OT (Okinawa Trough module), ME (Minami-Ensei Knoll site), Su (Sumisu Caldera site), IBMa (Izu-Bonin-Mariana arc module) and MT (Mariana Trough module). The OT, IBMa and MT are hereafter collectively referred to as the sub-regions of the Northwest Pacific while ME and Su are considered single-site outliers. The species nodes within a module represent species that are more closely associated to vent sites of that module than any other; many are not found at vent sites outside of their module (module endemics). Of the sub-region modules, the MT had the highest proportion of module endemics 80% (21/26), followed by OT with 57% (38/66), and finally IBMa with 47% (16/34) (Figure 5).

**Figure 5:**
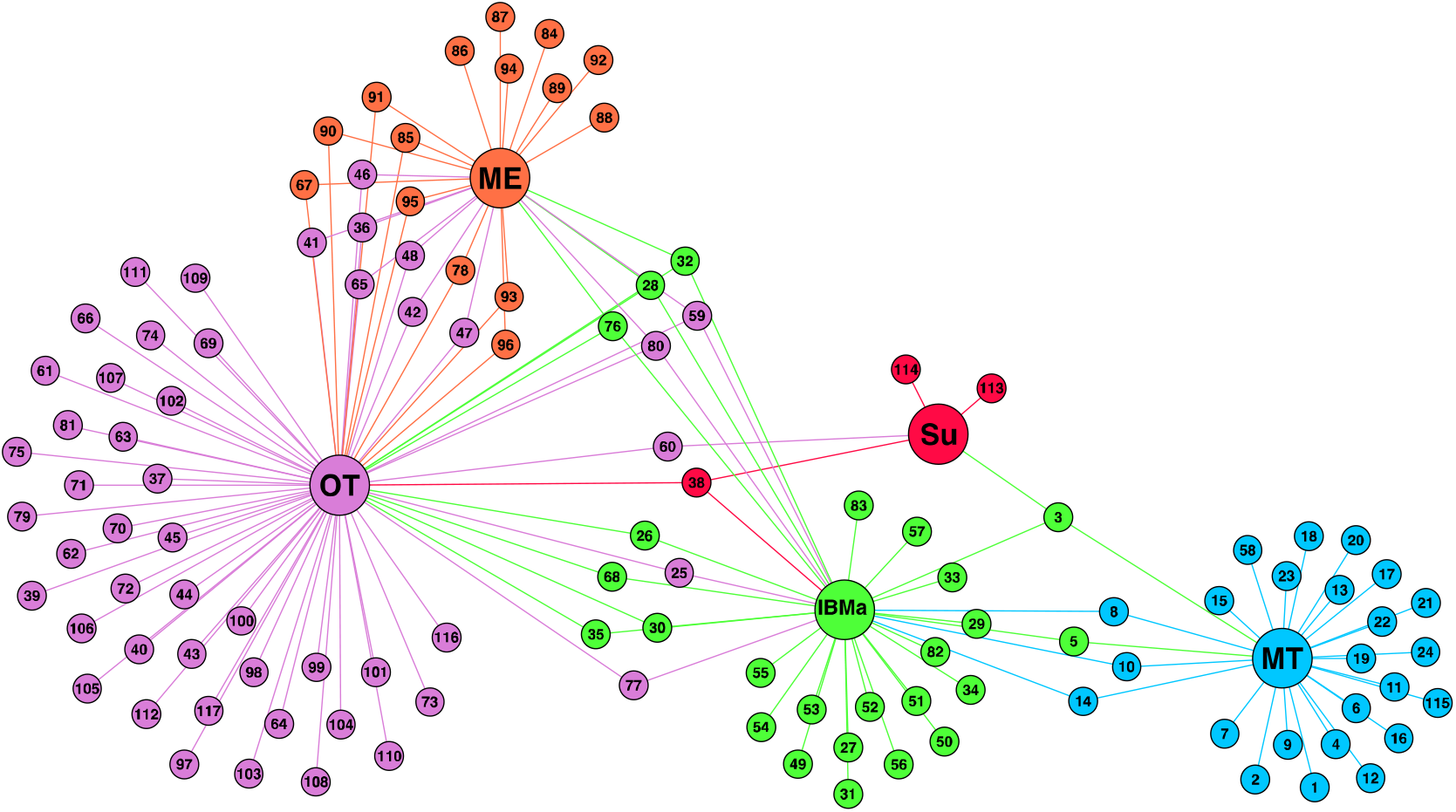
Bipartite network of Northwest Pacific with site nodes contracted by module group connected by species nodes. Node colours represent module groups; purple – Okinawa Trough (OT), orange – Minami-Ensei (ME), red – Sumisu (Su), green – Izu-Bonin-Mariana Arc (IBMa), blue– Mariana Trough (MT). Species nodes represent unique species, identified by number in supplemental table 2. While no species are shared between the three sub-regions, IBMa’s role as an intermediary between OT and MT is highlighted by its shared species.

Figure 6 presents the distribution of vent sites in terms of within-module degree (z_i_) and participation coefficient (P_i_) as well as the category of ‘universal roles’ (Guimerà and Amaral 2005) into which they fall (the same information for species nodes is summarised in supplemental table 2). The vent sites with the highest within-module degree of their respective sub-regions are Sakai (OT), Nikko (IBMa) and Alice Springs (MT). Of the three sub-regions, the vent site with the highest participation coefficient is Myojin Knoll of the IBMa (P_i_ =0.42). Other than the outliers, the five vent sites with the highest participation coefficients are Myojin Knoll, Northwest Rota and Myojinsho Caldera of the IBMa, and Daisan-Kume Knoll and North Knoll, Iheya Ridge of the OT module. All vent sites of the MT had relatively low participation coefficient, with Forecast having the highest 0.21). The vent sites that only contain module endemics have the lowest participation coefficients of 0 and are referred to as ‘ultra-periphery’ nodes. There are eleven such ultra-periphery nodes amongst the vent sites (Mokuyo Seamount, Chamorro volcano, Suiyo Seamount, Perseverance, Pika, Tarama Hill, East Diamante volcano, Snail, Daikoku volcano, Kasuga 2 Seamount, Nikko volcano), with representation from each of the three sub-regions.

**Figure 6:**
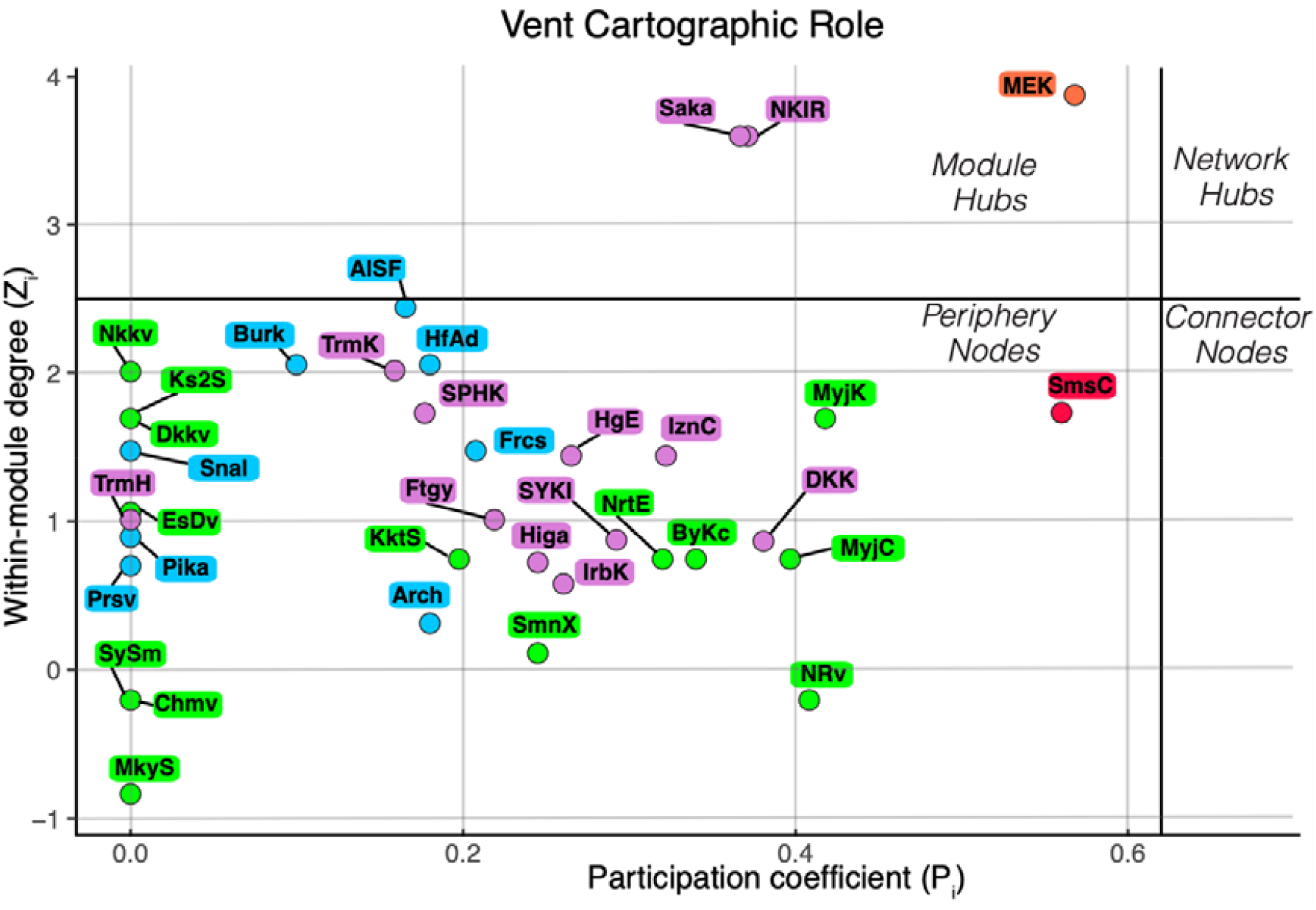
Within Module Degree and Participation Coefficient of each vent site in the bipartite network. Coloured by module membership as in figure 5; purple – OT, orange –ME, red –Su, green – IBMa, blue– MT.

The explanatory variables used in the redundancy analysis (broad-scale dbMEM, depth, and tectonic setting) were able to explain 92.8% of the module grouping, with tectonic setting and broad-scale dbMEM varying significantly between modules (p<.01 and .05 respectively).

## Discussion

The results obtained from the two networks generated in this study generally agree on the regional diversity structure as well as the significance of the environmental drivers we investigated. Although both networks are formed from species assemblage data, they each have distinct advantages in the way they can be interpreted. The similarity network builds upon common ecological analyses, but also extends from traditional clustering approaches to detect sites that act as intermediaries between clusters. Furthermore, displaying β-diversity as a network of similarity edges is arguably a much more intuitive way of visualising species diversity and inferred connectivity. The bipartite network is less intuitive in its presentation, but it can go beyond the detection of intermediary sites between clusters and identify sites that act as hubs of shared species within their cluster (module). Each of these two network methods have been previously employed in the study of various ecosystems at various scales, but not in combination. The discrete nature and dispersal processes that dictate connectivity at hydrothermal vents make them particularly suitable for the application of such network methods. The multivariate statistics applied to the species distribution data in this study found that the explanatory variables used (depth, tectonic setting and geodesic distance) were able to explain much of the variation between vent sites (Figure 4) and varied significantly between modules. Although these three explanatory variable were able to explain most of the variation, there are likely countless other abiotic, biotic and historical covariates that drive regional diversity at the biogeographic and metacommunity scale. The 10% of between-site variation explained by the environment (depth and tectonic setting) alone is likely due to unknown covariates. Tectonic setting, for example, can influence various parameters such as venting fluid composition, intensity and stability (Gamo et al. 2013; Mullineaux et al. 2018), which are considered drivers of community composition at vents (Juniper and Tunnicliffe 1997; Kojima and Watanabe 2015). Furthermore, differences in venting fluid composition between vent sites of the same tectonic setting, but different arc-back-arc basins, occur because of differences in the local sediment layers, material supply from the subducting slab and boiling points (Gamo et al. 2013). Although the vent-obligate species of this study are mostly independent of surface-derived food, depth is associated with the community structuring within this study as well as previous studies of regional hydrothermal vent communities in the Northwest Pacific (Nakajima et al. 2014; Kojima and Watanabe 2015; Watanabe and Kojima 2015; Watanabe et al. 2019). Important local controls on community composition can co-vary with depth, for example, the compositions and concentrations and concentrations of the chemicals in the vent fluid (Desbruyères et al. 2001). Depth often also co-varies with water-mass structure, which is a known biogeographic barrier within the Northwest Pacific (Brisbin et al. 2019). Depth is also known to have an influence on local vent fluid chemistry. For example, the depth of a vent site determines the boiling point of seawater and the boiling of venting fluid results in a phase separation that affects the local chemical composition of this venting fluid (Gamo et al. 2013). It is difficult to separate the influence of tectonic setting and depth in this region. Although arc and back-arc vent sites do overlap in terms of depth, particularly between Okinawa Trough and Izu-Bonin Arc, vent sites that occur on a volcanic arc are characteristically shallower than those that occur on the corresponding back-arc spreading centre (Figure 7). The two dbMEMs explain most of the species assemblage variation and represent their relative geographical position. The calculation of the fine-scale dbMEM assumes that vent sites in the Okinawa Trough are disconnected from the rest of the regional network while the broad-scale dbMEM assumes at least indirect connectivity between all vent sites in the Northwest Pacific. It is therefore unsurprising that the broad-scale dbMEM explains more variation between sites (22% compared to 7%) as it takes into account the species turnover among vent sites of the entire region. Connectivity among vent sites in the Northwest Pacific via larval dispersal has been shown to be possible under certain assumptions of larval dispersal behaviour by Mitarai *et al*. (2016). They also demonstrated that dispersal probability was variable among vent sites of the same sub-regional modules. It is possible that variable dispersal probabilities among vent sites in the Northwest Pacific is responsible for the variation explained by the fine- and broad-scale dbMEM variables. However, both dbMEMs also encompass all spatially autocorrelated variables that we were not able to account for in our analyses (Peres-Neto and Legendre, 2010). It is therefore likely that the variation explained by these spatial variables represents a dispersal limitation as well as spatially autocorrelated abiotic responses and biotic interactions among species (Thompson *et al*., 2020).

**Figure 7:**
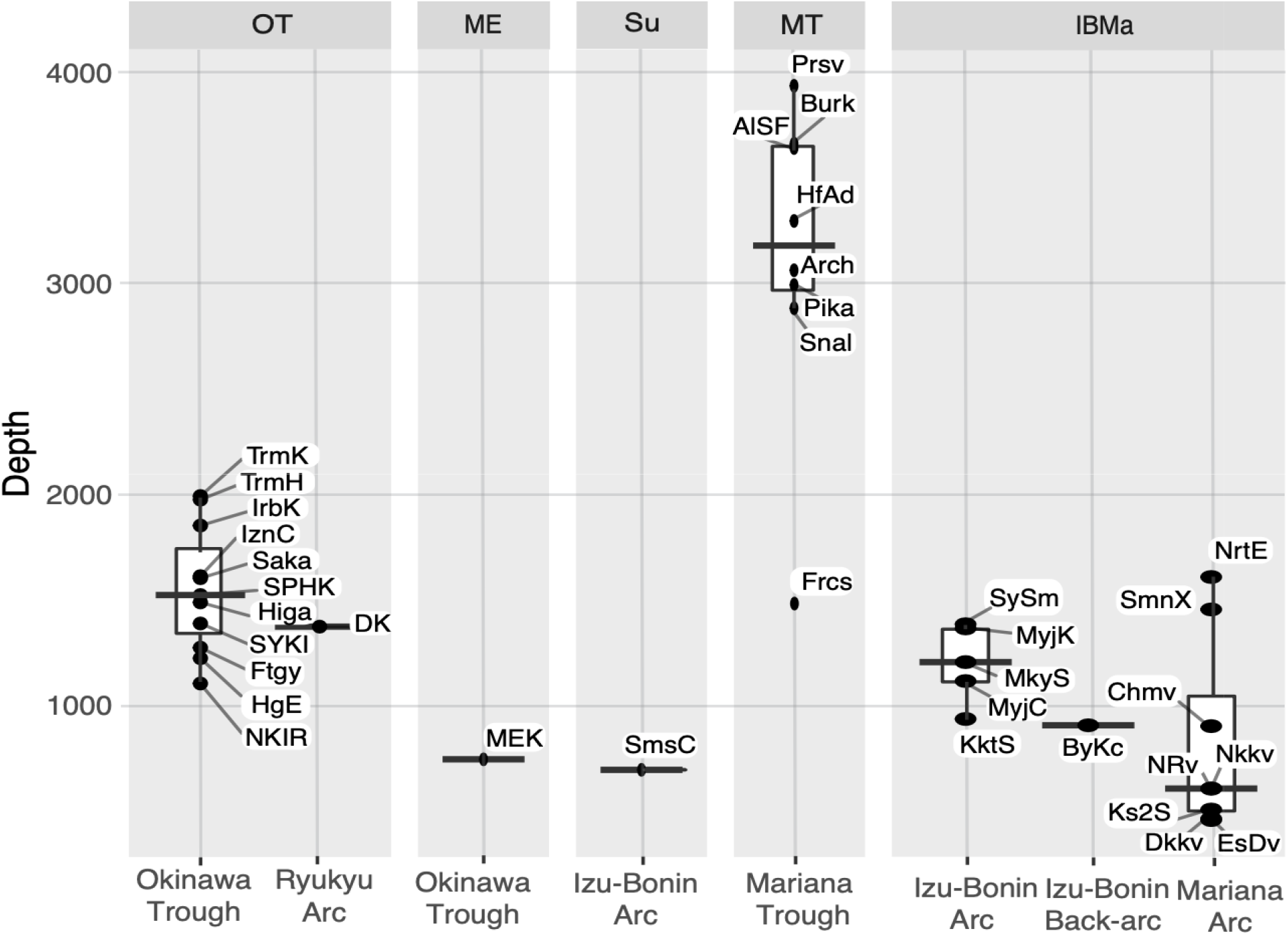
Depth distribution of vent sites separated into modules and subset into their basin

The uncertainty originating from the multivariate analyses and the lack of local biotic and abiotic parameters available to us precludes us from pinpointing the drivers of regional diversity. However, the significance of the parameters tested in combination with the structural aspects of the networks allow us to explore potential explanations.

### Regional Diversity Structure

At the scale of the entire region, the structure of the similarity network is defined as having “small world” properties due to the presence of tight clusters of nodes with key connectivity pathways (Figure 2) between them (Watts and Strogatz 1998). This same small world structure can be seen in the dispersal network of Mitarai *et al*. (2016) as well as from various marine metapopulation studies that evaluated the structure of dispersal networks (Kininmonth et al. 2010; Watson et al. 2011; Ramesh, Rising, and Oremus 2019). This small world structure was also found in the similarity network of Rozenfeld *et al*., (2008) who pioneered the application of network analysis to measures of pairwise site similarity values using the pairwise genetic similarities between populations of a seagrass species.

The clustering that gives the similarity network its small world properties is demonstrated by the arrangement of nodes based on their shared edges (Figure 2) and the high betweenness centrality of the key nodes that connect these three network clusters (Mariana Trough, Okinawa Trough and Izu-Bonin-Mariana Arc). This contrasts with the eleven SIMPROF clusters (Figure 3) that represent the lowest hierarchical level at which there are no significant structural differences between the vent site assemblages (Clarke *et* al., 2008). At the higher hierarchical level, it is possible to qualitatively divide the SIMPROF clusters into Mariana Trough, Okinawa Trough and Izu-Bonin-Mariana Arc ‘super-clusters’ in the same way as Kojima and Watanabe (2015). We detected the same three sub-regions using a modularity analysis of the bipartite network as well as the presence of two outlier sites (Minami-Ensei Knoll and Sumisu Caldera). At a glance, geographical proximity seems to be a strong driver of the module grouping (Figure 1.a) with OT to the west of the region, IBMa mostly to the northeast and MT to the southeast. It is likely that OT is spatially isolated based on the low dispersal probability connecting it to the rest of the region (Mitarai *et al*., 2016), but the differentiation between IBMa and MT can be better explained by the tectonic setting (Giguère and Tunnicliffe 2021) (Figure 2).

The occurrence of Sumisu Caldera as an outlier was detected across all methods. The similarity network disconnected this vent site from the network during the thresholding stage (Figure 2) and the SIMPROF results show a divergence of Sumisu Caldera from other vent sites in the Izu-Bonin-Mariana before their divergence from the Mariana Trough (Figure 3). Minami-Ensei Knoll however is not considered a clear outlier by the SIMPROF and similarity network methods. Sumisu Caldera and Minami-Ensei Knoll are the shallowest vent sites of the Izu-Bonin Arc and Okinawa Trough respectively (Figure 7). The occurrence of Sumisu as an outlier can be explained by its similarity to cold seeps in terms of its community composition (Nakajima et al. 2014) rather than its shallow depth, which is related to the fact that Sumisu Caldera is not very active for a vent and only exhibits weak diffuse flow (Iwabuchi, 1999). Minami-Ensei Knoll on the other hand does not resemble a cold seep in its community composition but does have a large proportion of endemic species and a few species uniquely shared with IBMa (Figure 5). It is possible that Minami-Ensei Knoll is distinct within the Okinawa Trough because of its shallow depth or high methane concentrations relative to the other vent sites in this area (recorded by Chiba (1993) and compared by Nakajima *et al*. (2014)).

There are minor discrepancies between the groups detected by the different methods but we will focus on those detected by the modularity analysis for the rest of the discussion. We have a preference for the results from the bipartite modularity analysis due to its objectivity compared to similarity network clusters and the additional insights it provides compared to the SIMPROF clusters (Bloomfield *et al*., 2018).

Furthermore, the significant difference of depth, tectonic setting and broad-scale dbMEM between the five modules from the MANOVA analysis validates the groups detected by the bipartite modularity analysis, which was not given information of these parameters *a-priori*.

### Inter-module Diversity Structure

The distinct sub-regions (excluding the two outlier vent sites) detected by the modularity analysis suggests the presence of biogeographic barriers within the Northwest Pacific, an area previously classified as a single biogeographic region (Bachraty et al. 2009). Previous studies that have applied this method to the analogous terrestrial ecosystem of island archipelagos (Mullineaux *et al*., 2018) have also inferred the presence of biogeographic barriers (Carstensen et al. 2012; Kougioumoutzis et al. 2014; Kougioumoutzis et al. 2017). Although we can infer that biogeographic barriers separate the three sub-regions we have detected, the proportion of shared species among them (in the case of OT and IBMa (Figure 5)) and their geographical overlap (in the case of IBMa and MT (Figure 1)) prohibits us from suggesting that the Northwest Pacific be further subdivided into additional biogeographic regions.

By contracting each module’s vent site nodes into a single node (Figure 5), it is clear to see which, and how many, species are shared between the distinct sub-regions. OT shares 12 species with IBMa and none with MT despite both MT and OT being predominantly composed of back-arc vent sites. The apparent role of IBMa as an intermediary of shared species between sub-regions (>50% species are shared with one of the other two sub-regions) indicates the importance of the spatial arrangement of these modules as IBMa vent sites are mostly found between those of the OT and MT in terms of oceanographic dispersal pathways (Mitarai et al. 2016). The role of the IBMa as a regional intermediary between OT and MT is also supported by the relatively high participation coefficient of many of their vent sites (Figure 6). Of the ten vent sites with the highest participation coefficient, four of them are in the Central/Northern Okinawa Trough and three of them (Bayonnaise Knoll caldera (ByKc), Myojin Knoll (MyjK) and Myojinsho Caldera (MyjC)) are from the northern Izu-Bonin Arc (the pathway of the Kuroshio current). Additionally, MyjK and MyjC have particularly high centrality scores due to their position along key pathways connecting the Okinawa Trough and the rest of the similarity network via Fuagoyama (Ftgy) and Daisan-Kume Knoll (DKK) (Figure 2). Daisan-Kume Knoll is the only Okinawa-Arc vent site in this study, so it shares both the tectonic setting as well as a similar depth with Myojin Knoll (Figure 7). However, the occurrence data for Daisan-Kume Knoll all come from the Gondou field on the western flank of the knoll, which is influenced by both arc and back-arc tectonic activity (Minami and Ohara 2017). The key pathway connecting the Izu-Bonin Arc to the Mariana Trough is via a Mariana Arc vent, Northwest Eifuku (NrtE), due to its connection to Forecast (Frcs) (Figure 2). The participation coefficient however, suggests that Northwest Rota-1 volcano (NRv) is a more important connection between sub-regions. This discrepancy is due to the emphasis the centrality measure ignoring ‘weak’ connections as they are removed by the thresholding step. The participation coefficient therefore suggests the importance of NRv due to its week connection to multiple vent sites of a different region while the centrality measure assumes these connections unimportant compared to a single ‘strong’ connection to the vent site Forecast (Frcs) of the Maraina Trough. The strong connection between NrtE and Frcs may be due to their depth proximity (Figure 7), while the weak connections of NRv and vent sites in the Mariana Trough may be driven by their geographic proximity (Figure 1). Previous studies have suggested that nodes with high centrality in vent similarity networks could have been historical stepping-stones between biogeographic provinces due to their geographic location or environmental features (Moalic *et al*., 2012; Kiel, 2016). Such historical biogeographic drivers have been found to have a strong role in present-day biogeography at vents (Kiel, 2017).

Although we were able to detect vent sites that act as important intermediaries among sub-regions, using the similarity network (Figure 2), the bipartite network was not able to detect equivalent ‘connector nodes’, based on Guimera and Amaral’s (2005b) universal cartographic roles (Figure 6). The lack of vent site connector nodes and prevalence of periphery nodes in the regional bipartite network is unsurprising considering the large number of module endemics (Figure 5). No species are present at all five modules or even all three sub-regions. Several species are present at three modules (two sub-regions and an outlier site). These species may have functional traits that allow them to cross these biogeographic barriers through strong dispersal or colonisation ability (Economo et al. 2015). The cartographic role of each species node can be found in supplemental table 2.

### Intra-module Diversity Structure

Only the bipartite modularity methods were we able to detect the relative importance of vent sites in driving within group diversity structure (Figure 6). Based on the shared species as well as geographic and environmental proximity, the sub-regions detected in this study may represent three distinct metacommunities of hydrothermal vent species in the region. Regardless of whether these are considered distinct metacommunities, biogeographic sub-regions or even regions, the guidance by Borthagaray et al. (2018) is that each identified module should be treated as independent biodiversity management units. This would suggest that the vent sites most important for maintaining network connectivity within their respective modules (highest z_i_ score) should be prioritised for protection to preserve regional diversity because they may also be key to maintaining gene flow within a metacommunity context. In this case, Sakai (Saka), Alice Springs Field (AlSF) and Nikko volcano (Nkkv) should be prioritised for protection in their respective sub-regions. Special consideration for conservation should be given to the outlier vent sites of Sumisu and Minami-Ensei Knoll due to their high proportion of endemic species (Figure 5). In their study of bird communities on island archipelagos, Cartensen *et al*., (2012) used the same bipartite network methods to conclude that module hub islands are sources of species for periphery node islands. This conclusion was partly based on the module hub islands being consistently larger than periphery node islands, a feature we are unable to confirm for hydrothermal vent sites in our study.

There is no evidence for immediate anthropogenic risks, in the form of deep-sea mining, to the vent sites of the Izu-Bonin-Mariana Arc and Mariana Back-arc, some of which fall within the US Marine National Monument of the Marianas (Menini and Van Dover 2019). However, the central Okinawa Trough is confirmed as an area of interest for planned hydrothermal vent mining (Okamoto, Shiokawa, and Kawano 2019), although specific vent sites have not been publicly confirmed as targets for mining. Those vent sites in the Okinawa Trough that are of particular interests in terms of their seafloor massive sulfide deposits have been identified as Sakai, Izena Cauldron, and Daisan-Kume Knoll (Ishibashi et al. 2015; Minami and Ohara 2017). As a ‘module hub’ for the OT sub-region, with the highest within module degree, any disturbance to Sakai from mining activity will have a disproportionately strong impact on the linkages among vent sites within the Okinawa Trough metacommunity. The high betweenness centrality of Daisan-Kume Knoll suggests it plays an important role in sharing species between the Okinawa Trough and Izu-Bonin Arc, this may be due to its tectonic features which make it an intermediary in terms of local environmental conditions.

## Conclusion

We conclude that the Northwest Pacific hydrothermal vent bioregion is divided into three distinct sub-regions - Okinawa Trough, Izu-Bonin-Mariana Arc and Mariana Back-arc - that do share species but have distinct metacommunities separated by biogeographic barriers. The Minami-Ensei Knoll and Sumisu vent sites contain species assemblages distinct from all three of the sub-regions and should be considered outliers that indicate the importance of local environmental factors in distinguishing vent site assemblage structure. Most of the variation in species composition between individual vent sites and between the sub-regions can be explained by local environmental filters associated with tectonic setting and depth, as well as the geodesic distance between vent sites.

The shared species and linkages across the distinct sub-regions of the Northwest Pacific can be disproportionately attributed to a small number of vent sites, namely Daisan-Kume Knoll, Myojin Knoll, Northwest Eifuku, and Forecast. Disturbances to these nodes with high centrality would have relatively strong impacts on maintaining connectivity between the three sub-regions, as they are key nodes in preventing the collapse of the regional network (Thompson et al., 2015). The vent sites most important for linking within the sub-regions of the Okinawa Trough, Izu-Bonin-Mariana Arc, and Mariana Back-arc are Sakai, Nikko Volcano, and Alice Springs Field, respectively. These three vent sites play the most important role in maintaining biodiversity in the Northwest Pacific on time-scales pertinent to conservation, by connecting the sub-regional metacommunities through shared species.

